# Intracortical BCI Performance is Robust to Changes in Attentional Load During Dual-Tasking

**DOI:** 10.64898/2026.06.16.732398

**Authors:** E. Canario, C. Shearer, M. Akcakaya, D. Weber, S. Chase, J. L. Collinger

## Abstract

High performance intracortical brain-computer interface (iBCI) control has been demonstrated in research settings, but performance can still vary within and between sessions. One potential source of this variability is the change in attentional load that comes from processing naturally occurring distractors such as thoughts, sounds, fatigue, or pain. To improve the consistency of iBCI performance in real-world environments where this sort of multi-tasking is inevitable, we must understand how shifts in attention can impact performance. Here we examined the effect of attentional load on iBCI performance and movement-related neural activity using a 2D cursor translation + click iBCI task paired with an N-Back working memory task to increase attentional load during dual-task performance. Two participants (P2 and P4) with tetraplegia completed the study while enrolled in a long-term clinical trial of an iBCI device (NCT1894802). Common neural correlates of attention (theta and alpha band power) were measured with simultaneously recorded scalp electroencephalography (EEG). While the EEG recordings and difficulty ratings suggested increased attentional load during dual tasking, iBCI performance was quite robust across the various dual tasking conditions. One participant, P2, experienced a small but significant increase in trial completion time and normalized path length during the mild attentional load condition. Signal quality differences between the two participants may have impacted the results, as P2 had lower signal quality and was therefore likely more vulnerable to attentional load. P4’s higher signal quality likely allowed him to accommodate increased attentional load without a drop in performance. Overall, iBCI performance appears to be robust to attentional load, but the complex trends observed here reflect a need for continued investigation of BCI use under different cognitive states to elucidate potential challenges and compensatory mechanisms across participants.

## INTRODUCTION

Intracortical BCIs have achieved a high level of performance in controlled laboratory environments (Card et al., 2024; Collinger, Wodlinger, et al., 2013; Pandarinath et al., 2017). This level of performance has motivated a transition to real-world testing in home environments in both academic and industry-led clinical trials (Card et al., 2025; Neuralink, 2026 Oxley, 2025; Simeral et al., 2021; Weiss et al., 2019). However, assistive technologies are often abandoned by users due to reasons such as mismatched user needs and device function, difficulty of use, and poor performance (Phillips & Zhao, 1993). Potential BCI users desire systems that are highly reliable and require minimal setup time (Blabe et al., 2015; Collinger, Boninger, et al., 2013; Huggins et al., 2011). Reliable and stable BCI control can be challenging to achieve because neural recordings are inherently unstable due to physical factors such as micro-motion of arrays (Perge et al., 2013) or context-dependent changes in the underlying neural population activity (Perge et al., 2013; Sussillo et al., 2016; Tadipatri et al., 2017). Some progress has been made using decoding algorithms that rely on neural networks to enable stable control with little to no recalibration (Card et al., 2024; Jude et al., 2025) but their robustness to changes in mental state or environment has not been fully evaluated.

In other motor control contexts, attention is known to impact task performance (Draheim et al., 2022; Engström et al., 2017; Pashler, 1994), but this has been less investigated in BCI. Furthermore, the few studies of attention focus primarily on BCIs that rely on scalp electroencephalography (EEG), with few examining the effect of changes in attention on implanted BCI control. EEG BCIs record neural activity with electrodes placed on the scalp. There are several types of EEG BCIs, with most being based on evoked potentials or motor imagery. Evoked potentials, such as the P300 or the steady-state visually evoked potential (SSVEP), are time-locked neural responses that occur in response to a stimulus (İşcan & Nikulin, 2018; McFarland et al., 2011). A P300-based BCI speller was found to have lower performance in high mental workload conditions as well as when the user became more fatigued (Käthner et al., 2014). Another study found overall classification performance dropped when using an SSVEP based BCI during dual-tasking (İşcan & Nikulin, 2018) though these results depended on the context (speech vs listening vs thinking) and level of cognitive load. Motor imagery-based BCIs rely on patterns of neural activity, such as sensorimotor rhythm desynchronization, that occur during imagined movement and enable more continuous control dimensions than evoked potentials (McFarland & Wolpaw, 2017). In an EEG-BCI cursor control task, subjects were found to be sensitive to speech-related distractors, which decreased their target acquisition and path efficiency (Foldes & Taylor, 2013). Another EEG-BCI study found that distractors of various cognitive loads impacted lower-performing participants but not higher-performing ones (Emami & Chau, 2020). Indeed, subject state often has a very complicated relationship with performance that is dependent on multiple factors. For example, one study found that fatigue, frustration, and attention could have either positive or negative effects on EEG-BCI performance (Myrden & Chau, 2015). Specifically, they noted that performance was best when fatigue was moderate and frustration was high. They hypothesized that fatigue was a result of increased effort and frustration served as a motivating factor to improve performance. Taken together, these studies demonstrate that an interplay of different factors such as fatigue, attention, frustration, and BCI proficiency can result in varied outcomes for overall performance.

Attention has been less studied for intracortical BCIs, but some evidence suggests that intracortical BCIs may be more robust to distractors than EEG-based BCIs, with only minimal performance loss during simple motor tasks performed with various distractions (Guthrie et al., 2021). However, a limitation of this previous study is that the BCI task was relatively simple, as participants only had to move between two nearby targets repeatedly, so it is unclear how attention may impact iBCI use particularly for more complex and realistic tasks. Evidence from attentional studies in other contexts such as driving finds that tasks that are more difficult, requiring active thought and increased effort, are more affected by distractors (Engström et al., 2017). One intracortical BCI study found that cursor control completion times increased during continuous speech but not shorter forms of speech (Stavisky et al., 2020). This suggests that more involved distractor tasks have a greater effect on BCI control.

Beyond just performance, cognitive factors can also directly impact the neural activity that drives movement. For example, distractors can disrupt the sensorimotor rhythm desynchronization often used to drive EEG-BCIs, even if there is no effect on performance (Emami & Chau, 2018). Arousal can modulate brain-wide neural activity (McCormick et al., 2020; Musall et al., 2019; Stringer et al., 2019) including motor activity during BCI task learning (Hennig et al., 2021). Further, motor cortical activity can change depending on expected reward or motivational state (An et al., 2019; Ramakrishnan et al., 2017; Ramkumar et al., 2016; Smoulder et al., 2024).

To better understand how motor-related neural signals and BCI performance change with attentional load, we recorded scalp EEG during iBCI use. Neural correlates of attention are commonly measured with EEG during concurrent motor tasks. For example, EEG recordings showed increased frontal area theta band power and decreased parietal area alpha band power while participants were driving under distracting conditions (Wang et al., 2018). Similar trends were observed during dual task performance involving a motor and working memory (N-back) experiment (Ozdemir et al., 2016) and during a dual task standing and verbal math experiment (Kahya et al., 2022). Frontal theta power (∼4-8 Hz) is thought to be involved in executive function during cognitive tasks (Cavanagh & Frank, 2014). Alpha power (∼8-12 Hz) in the parietal-occipital region is thought to be involved in the allocation of cortical resources to internal or external tasks (Magosso et al., 2019).

Here we aimed to investigate the impact of attentional load on iBCI performance and the underlying movement-related neural activity in two human intracortical BCI users. Participants used the iBCI to perform a complex cursor control task involving click and drag actions (Dekleva et al., 2021) combined with the N-Back working memory task (Kirchner, 1958) to increase attentional load, emulating real-world distractions. The N-Back task impacted attention but had little effect on BCI performance. Subject-specific effects were observed in the motor-related neural activity, though the changes were fairly small in magnitude, which may suggest that each had a different compensatory strategy based on their baseline BCI performance. One participant had been implanted for a much longer duration and therefore had lower signal recording quality and baseline BCI performance. Despite these differences, iBCI cursor control was generally robust to attentional load. We demonstrated the feasibility of concurrent EEG recording and iBCI control, which may enable future studies that wish to examine how iBCIs generalize to different settings, where the user may experience stressors, such as pain or fatigue, that impact their ability to attend to BCI use.

## Materials and Methods

### Data Collection

Two participants, P2 (28 years old at implant, C5 ASIA B spinal cord injury) and P4 (31 years old at implant, C4 ASIA A spinal cord injury), with tetraplegia who were participating in a clinical trial of an iBCI device for restoring upper limb function completed the experiments for this project. This study was conducted under an FDA Investigational Device Exemption (NCT1894802) in accordance with the principles embodied in the Declaration of Helsinki and approved by the University of Pittsburgh Institutional Review Board. Informed consent was obtained prior to any experimental procedures. Study participants each had two intracortical microelectrode arrays (NeuroPort Electrodes, Blackrock Neurotech, Salt Lake City, UT) implanted in the motor cortex (88 channels each for P2 and 96 for P4) and two 32 channel arrays in the somatosensory cortex. P2 was implanted approximately 8 years prior to data collection, while P4 was implanted approximately 6 months prior to data collection.

Intracortical signals were collected at 30 kHz using the NeuroPort System (Blackrock Neurotech) and filtered with a 4^th^ order 250 Hz high-pass filter. Threshold crossing events (-4.5 RMS) were recorded and binned (every 20 ms) to estimate spike rates. The binned spike counts were then smoothed with an exponential smoothing function with a 440 ms window. To isolate local field potentials (LFPs), the 30 kHz data were low-pass filtered with a 4^th^ order filter and then down-sampled to 1 kHz. We calculated the mean channel raw beta band frequency power (13-30 Hz) from each trial. This was done by first calculating a short-time Fourier transform for each trial and then averaging across time within a specific beta band. A few trials (3 for P2, 2 for P4) were rejected due to software error resulting in no intracortical data. Figure 1A displays the approximate location of arrays (represented by blue squares) and percutaneous pedestal connectors (represented by the blue circles).

**Figure 1:**
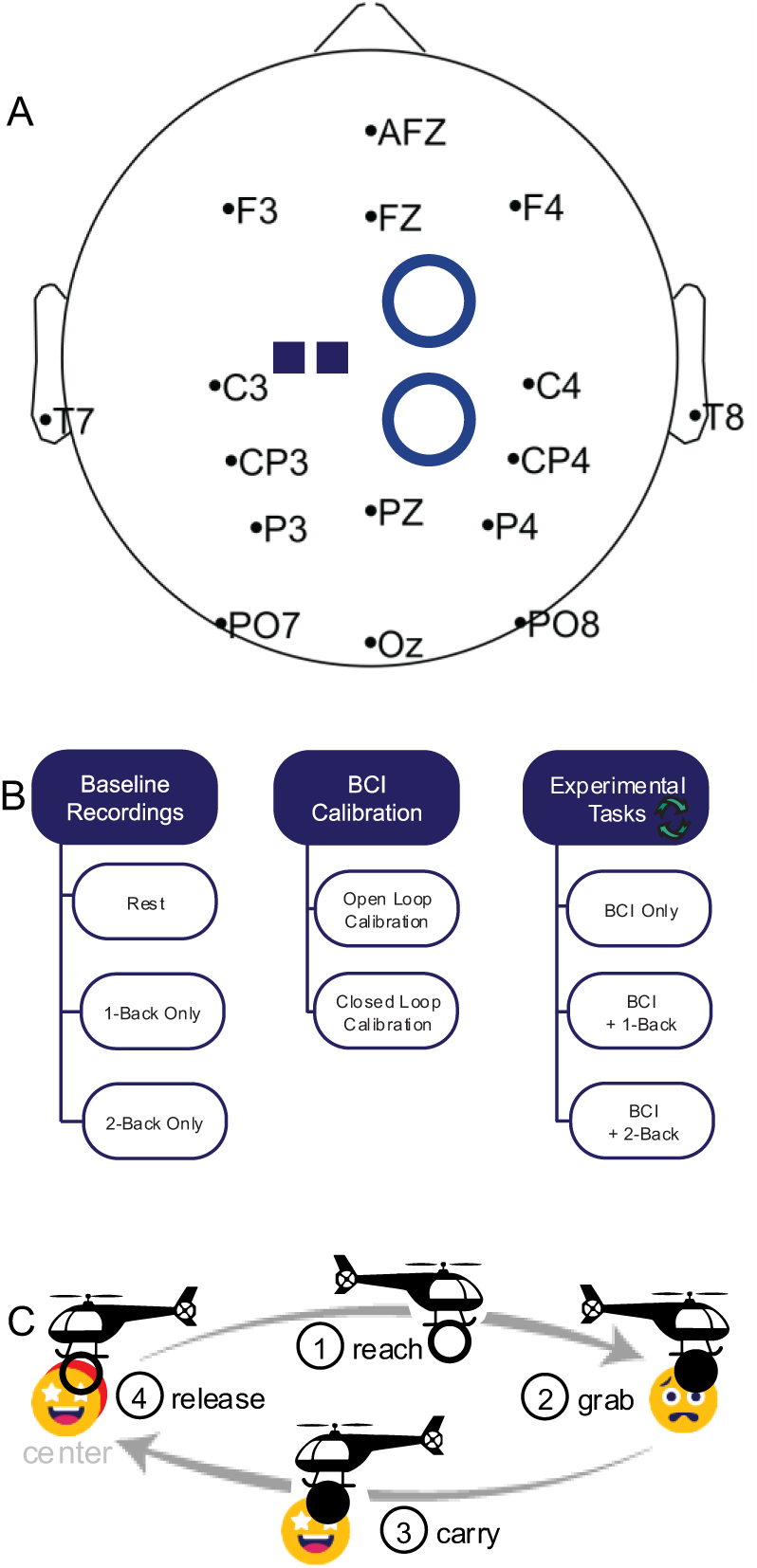
A: Location and names of the 16 EEG channels. Blue ovals indicate the approximate location of percutaneous pedestal connectors and blue squares indicate the approximate location of intracortical motor arrays on the brain. Frontal electrodes are marked by the letters F and AFZ. Parietal-occipital electrodes are labeled with the letter P, PO, or O. Central electrodes are labeled with C or CP, and the T label indicates electrodes on the temporal region. B: Each session consisted of baseline recordings, BCI calibration, and then BCI control with and without N-back. Baseline recordings included three minutes of rest, 1-Back only, and 2-Back only. The dual-tasking experimental conditions were block randomized each session. C: BCI task phases. First, during the reach phase the participant navigated the cursor (helicopter) to a target (face emoji). Next, they clicked to grab the target and carry it back to center where they ‘unclicked’ to release the target. Trials were separated by a 2-second intertrial interval.

To provide a more global measurement of neural activity, EEG data were collected using 16 g.tec ladybird active electrodes and a g.USBamp biosignal amplifier (g.tec medical engineering, Schiedlberg, Austria) at 256 Hz. The right mastoid was used as the reference electrode, and the left mastoid was used as ground. Channel locations are shown in Figure 1A derived from the international 10-20 system of EEG electrode placement (Jasper, 1961) with slight adjustments to accommodate the percutaneous pedestals that allow connection to the implanted intracortical electrodes. EEG data were recorded in BCI2000 (Schalk et al., 2004) and processed in EEGLAB (Delorme & Makeig, 2004). EEG data were filtered with a 0.5-55 Hz bandpass and then bad channels were visually identified and removed. At most, 1-2 channels were removed per set. Next, we used artifact subspace reconstruction (Mullen et al., 2015) with a standard deviation of 15 to automatically identify bad data segments and remove them. We then performed a common average reference with all remaining channels. Finally, ICA was performed to remove eye-related components that might contaminate the data. For data collected while the participant was at rest, this pipeline resulted in 5% (P2) or 18% (P4) of the data being rejected. To avoid discontinuities, individual trials were rejected if they contained any rejected data segments, leading to up to 3.5% (P2) or 41.5% (P4) of trials being rejected. More trials were rejected for P4 due to greater noise for reasons such as more movement artifacts and worse electrode contact with the scalp. Table 1 indicates the total number of EEG trials per condition per participant after rejection.

**Table 1:** Number of EEG trials included for analysis. Excluded trials were rejected due to artifact contamination.

| Participant | BCI Only Trials | BCI+1-Back Trials | BCI+2-Back Trials |
| --- | --- | --- | --- |
| P2 | 143/144 (99%) | 144/147 (97%) | 153/160 (95%) |
| P4 | 152/232 (65%) | 118/208 (56%) | 109/208 (52%) |

### Session Design

Figure 1B shows the overall session design that included baseline recordings, BCI calibration, and experimental dual tasking conditions involving BCI and the N-back working memory test (Kirchner, 1958). Participants were familiarized with the N-Back through practice trials before any experimental conditions were recorded. The N-Back is a task designed to test working memory. In the N-Back conditions, participants were presented with letters via audio with an inter-trial interval of 2.5 seconds while they performed the BCI task. Participants were instructed to verbally indicate when the letter presented matched the letter presented N (where N=1 or 2) letters ago. The presentation of letters was randomized. An example of 2-Back would be: I G K **G** B L, where the subject would say “match” on the bolded letter, G. As with many dual task studies, we expected that managing this secondary task required more attentional resources from participants due to the greater attentional load compared to only doing one task.

At the beginning of each session, we recorded EEG and intracortical data during a three-minute resting baseline period. Participants were instructed to look at a fixation cross while minimizing movement. We also collected EEG and intracortical data during performance of the 1-Back and 2-Back conditions without any BCI control, each for 3 minutes. We then calibrated the BCI decoder and proceeded to the experimental conditions (See *Task Design* for details). EEG and intracortical neural data were recorded while the participants performed one of three conditions: 1) BCI Only, 2) BCI with mild attentional load (BCI+1-Back), and (3) BCI with moderate attentional load (BCI+2-Back). Participants were told to focus on the BCI cursor task as the primary task but to try their best to complete the N-Back accurately. The condition order was block randomized each session to minimize order effects. We collected 1-2 sets of 16-24 trials of each condition depending on time constraints (Table 1). After each condition, participants were asked to rate their mental effort on a scale from 1-10 (Paas, 1992). We collected 5 sessions with P2 and 6 with P4.

### Task Design

To calibrate the BCI decoder, we began with open-loop calibration where the participants attempted to perform a center-out-and-back cursor control task in both clicked and unclicked states (Dekleva et al., 2021). Neural data was recorded while the cursor movement was controlled by the computer. Factor analysis, a type of dimensionality reduction, was used to produce a 20-dimensional neural state space. This state space was used to train a Kalman filter to decode 2D cursor velocity and two LDA classifiers to decode click and unclick. We repeated the process with a partially assisted closed-loop calibration in which off-target movements were attenuated to train a second decoder that was then used for the remainder of the session.

Subjects performed an iBCI-controlled cursor task, which was a gamified center-out grasp and carry task (Figure 1C). This 2D+click BCI task was designed to emulate mouse use on a computer. It consisted of an initial presentation phase where the target is presented, a reach phase where participants reach towards the target, a click phase where they click the target, and a center phase where they return to the center to end the trial. A black screen was then shown for two seconds to serve as an intertrial rest period. Participants had a maximum of 20 seconds to complete each non-presentation phase, where failure in any phase meant failing the trial. A typical block consisted of 8 trials, with 2 or 3 blocks per set. The order of the sets was randomized for each session.

BCI task performance measures included success rate (per block of 8 trials) and reach phase completion time, total trial completion time, and normalized path length for the successful trials. Normalized path length was calculated as the ratio of the actual cursor path length to the ideal cursor path length (where a straight line to the target is equal to 1) during the reach phase. Total trial completion time was included to assess impact on the full trial with translation and clicking, while reach phase completion time focuses on the simpler translation period.

### Neural Data Analysis

We extracted the attention-related measures of EEG band power for the theta (4-7 Hz) and alpha (8-12 Hz) bands as well as the sensorimotor rhythm (beta band, 13-30 Hz). We first calculated the event-related spectral perturbation (ERSP) for every trial using EEGLAB, and computed the average power in the theta, alpha, and beta bands for the first two seconds of the reach phase, obtaining a trial level measure that captured the initial period of movement towards a target before participants began self-correcting in response to error. EEG features of attention are present broadly in the brain but are strongest in certain regions, such as alpha primarily appearing in the parietal region and theta in the frontal (Emami & Chau, 2020). We used a region of interest approach and averaged power in the parietal-occipital channels (PO7, PO8, P4, P3, Pz) to obtain alpha power and frontal area channels (AFz, Fz, F3, and F4) to obtain theta power as is common in studies of attention (Scharinger et al., 2017). Beta band power is expected to decrease during movement periods so a smaller value of beta band power during the reach phase would indicate stronger movement-related activity. For examples of the EEG signal, see Figure S1 in the Supplementary Material. From the intracortical spike rate data, we calculated the mean firing rate in each trial (consisting of the first two seconds after reach begins, as in the EEG processing) for all conditions (Wilson et al., 2023). We also computed the neural reaction time based on the time it took to reach the peak firing rate during the first two seconds of the trial’s reach phase. We also computed beta band power from the intracortical local field potential (LFP) recordings. We then calculated the ERSP for every trial across the first two seconds of reach. We averaged this across all motor channels (176 for P2, 192 for P4). Finally, beta band power was then averaged across frequencies and time to obtain a trial level metric of the LFP beta band power.

### Statistical Testing

Data were combined across days for statistical testing. Normalized path length, total trial and reach trial completion times were found to have skewed distributions so they were log-transformed to aid normality. For every metric of performance, attention, and motor signal, we ran separate statistical tests. First, we ran a regression model to identify the effect of condition of outcome metrics, with session as a random effect. Then, we assessed the significance of the model with an ANOVA, and finally, performed post-hoc comparisons using the Tukey Method to compare conditions.

## RESULTS

### iBCI Performance is Stable under Increasing Attentional Load

iBCI performance was generally robust to attentional load challenges induced by the dual tasking N-back paradigm (Figure 2). Neither participant showed significant differences in success rate (P2: F = 0.848, p = 0.435; P4: F = 1.676, p = 0.194) or reach phase completion time (P2: F = 2.701, p = 0.068; P4: F = 1.307, p = 0.271). P2 had higher total trial completion time in BCI + 1-Back (median: 11.63 seconds, IQR: 8.32-19.60) compared to BCI + 2-Back (median: 10.42 seconds, IQR: 7.76-13.50, p < 0.05), but P4 exhibited no differences (P2: F = 5.751, p = 0.003; P4: F = 0.405, p = 0.667). Additionally, P2 also had an increase in trial normalized path length in BCI+1-Back (median: 0.16 A.U., IQR: 0.07-0.28) as compared to BCI+2-Back (median: 0.13 a.u., IQR: 0.08-0.21, p < 0.05) but this was not the case in P4 who had no significant differences (P2: F = 4.739, p = 0.009; P4: F = 0.793, p = 0.453). Dual tasking was rated as slightly more difficult than BCI control alone (Figure S2, Supplementary Material).

**Figure 2:**
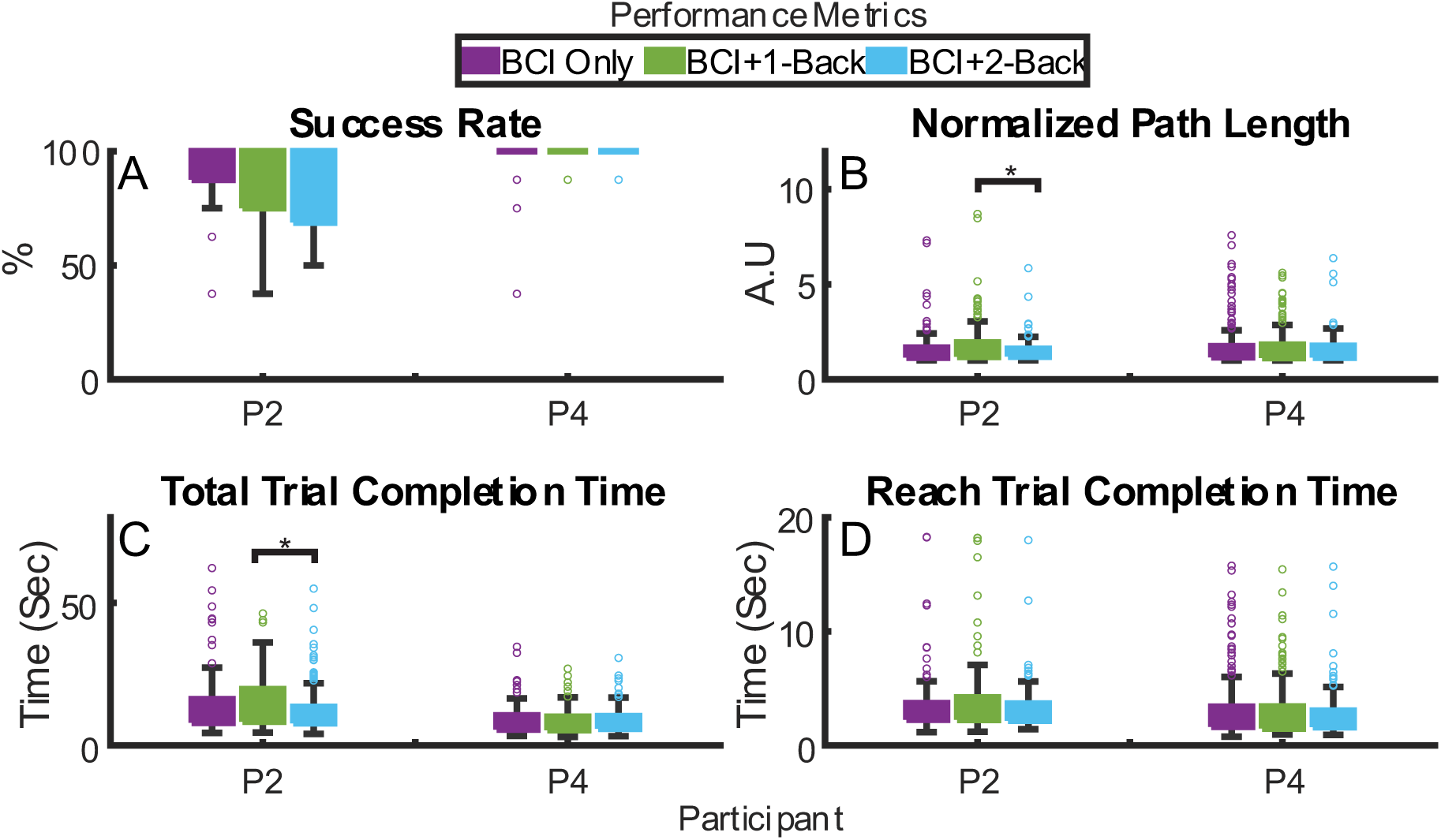
Performance metrics during dual tasking conditions. A: Success Rate, B: Normalized Path Length, C: Total Trial Completion Time, D: Reach Trial Completion Time. P2 exhibited a higher path length and total trial completion time in BCI + 1-Back compared to BCI + 2-Back, indicating worse performance.

### Neural Correlates of Attention are Modulated by Attentional Load

EEG metrics of attention were well modulated by performance of the N-Back task (without BCI control) as compared to rest. Theta power was significantly different among the three baseline conditions (rest, 1-back, & 2-back) for P2 and P4 (Figure 3; P2: F = 168.08, p < 0.001; P4: F = 22.784, p < 0.001), with theta power being the highest for the 1-Back condition (p < 0.05). For P2, theta power was also higher during the 2-back condition as compared to rest (p < 0.05), but to a lesser degree than during the 1-back. Alpha power was also higher during the 1-back condition as compared to rest and BCI + 2-Back in P2, while in P4 it was higher in both of the N-back only conditions compared to rest (P2: F = 165.11, p < 0.001; P4: F = 23.539, p < 0.001). This result is in the opposite direction from what was expected, as we initially hypothesized there would be a linear decrease in response to increasing attentional load. Thus, both alpha and theta (for both participants) seem to increase the most in 1-Back but then decrease in 2-Back compared to 1-Back and/or rest.

**Figure 3:**
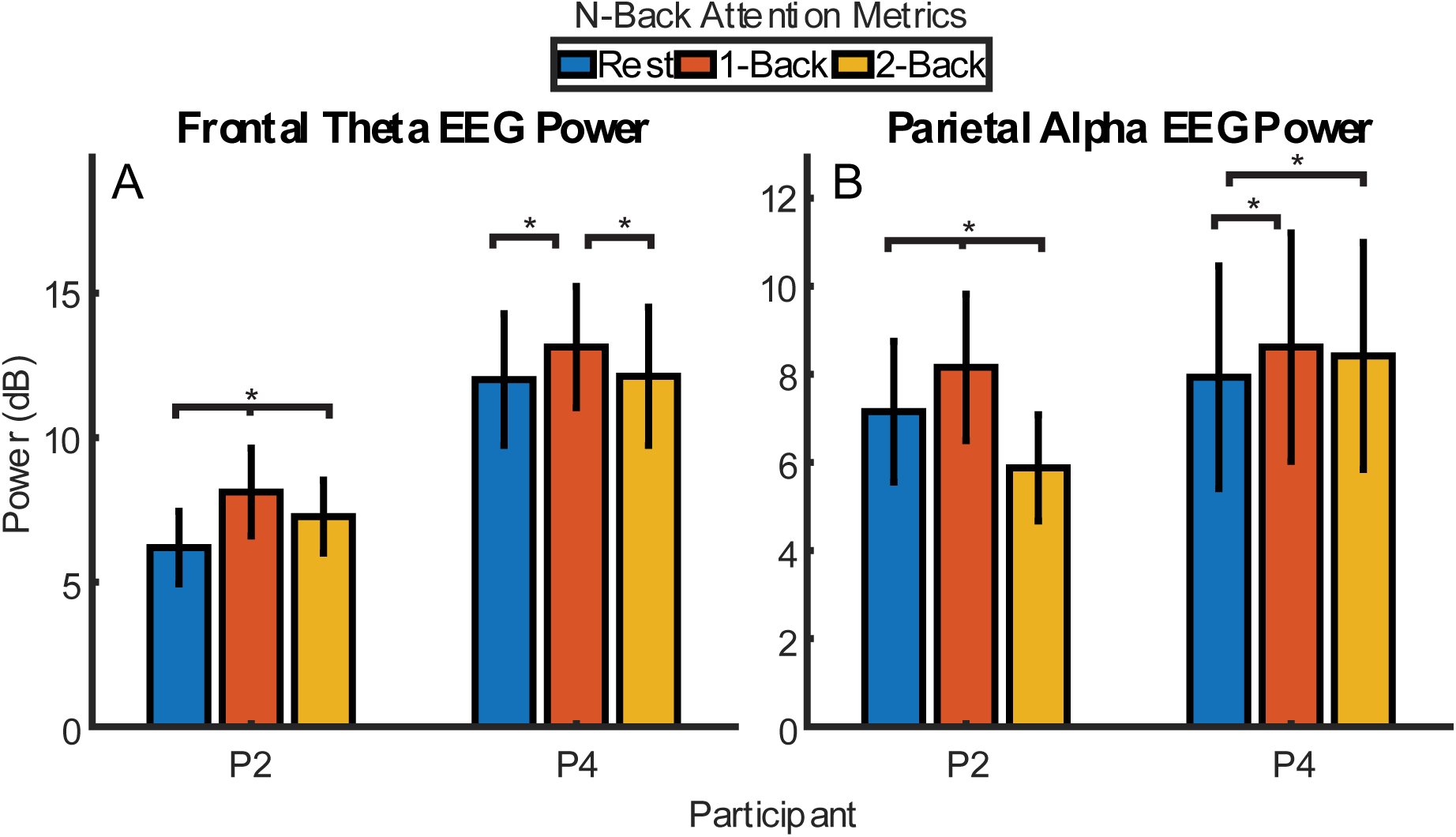
Comparing N-Back only to resting state. A Frontal Theta EEG Power, B Parietal Alpha EEG Power. Participants displayed higher theta power in 1-Back compared to rest, with P2 also having higher theta power in 2-Back. Participants displayed higher alpha power in 1-Back compared to rest, with P2’s 2-Back alpha power being lower than other conditions.

In dual-tasking (BCI + N-back), we see similar trends. Specifically, when looking at neural correlates of attention (Figure 4), we see changes in frontal theta power for both participants across dual-tasking conditions (P2: F = 6.243, p = 0.002; P4: F = 4.609, p = 0.011). For P2, theta power is higher in the BCI+1-Back compared to BCI Only (p < 0.05) and BCI + 2-Back (p < 0.05). For P4, theta power is higher in the BCI+2-Back condition (p < 0.05) as compared to BCI Only. When looking at alpha power in the parietal region, we found a significant difference across conditions for P2 but not P4 (P2: F = 10.271, p < 0.001; P4: F = 0.6744, p = 0.523). For P2, alpha power was higher in the BCI+1-Back condition compared to BCI Only (p < 0.05) and BCI + 2-Back (p < 0.05).

**Figure 4:**
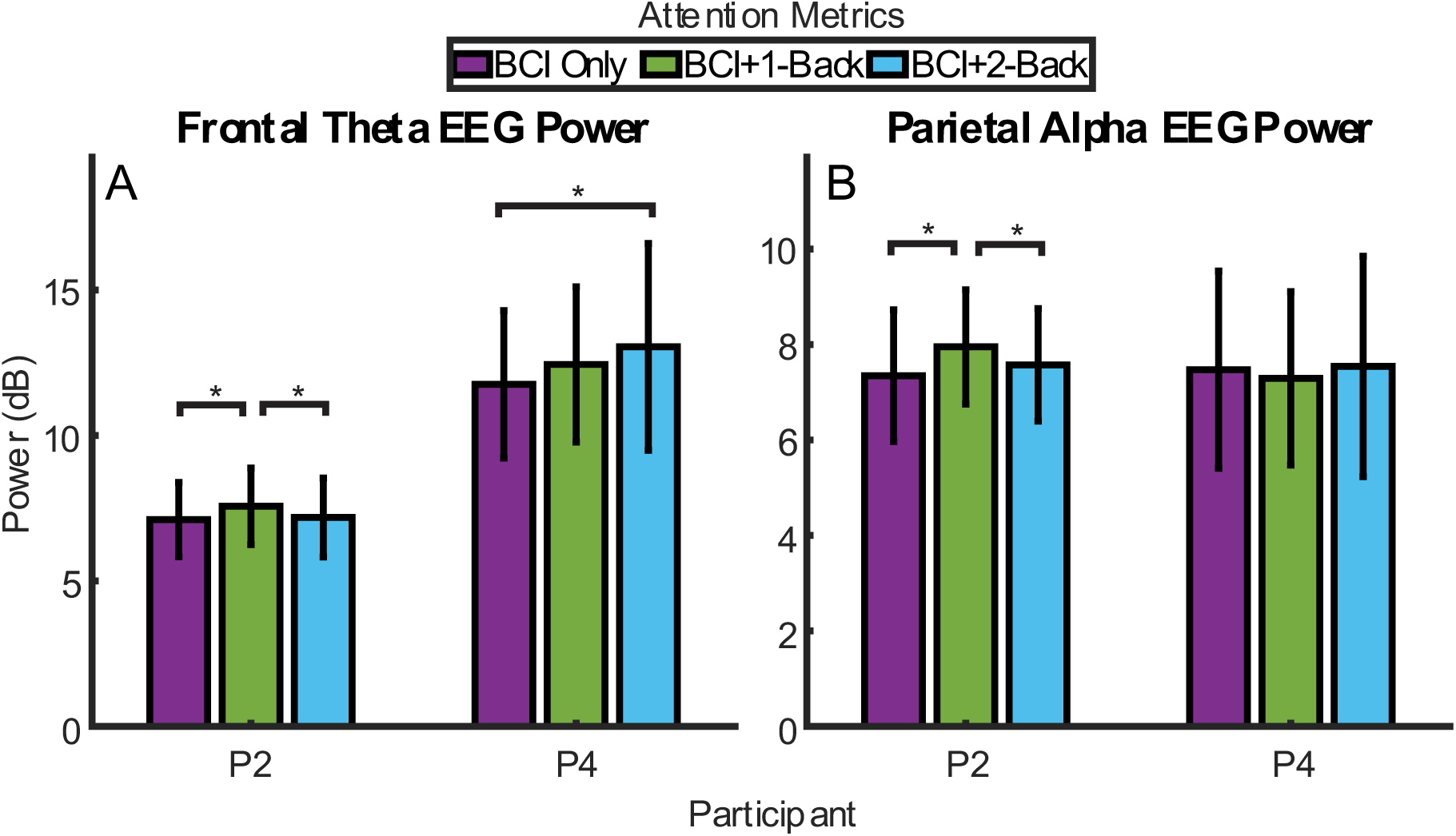
Comparing dual tasking to BCI Only. A: Frontal Theta EEG Power, B Parietal Alpha EEG Power. Participants displayed higher theta power in either BCI+1-Back (P2) or BCI+2-Back (P4) compared to BCI Only. P2 displayed higher alpha power in BCI+1-Back compared to BCI Only.

### Subject-specific Changes in Movement-related Signals Under Attentional Load

We analyzed three movement-related signals for both participants to examine whether or not attentional load impacts the high-level intention signal in motor cortex. We found that movement-related activity was generally consistent across conditions (Figure 5), mirroring the trends in BCI performance (Figure 4). Neither participant showed differences in the EEG beta band power (P2: F = 1.660, p = 0.191; P4: F = 2.066, p = 0.128). Only P2 demonstrated any difference in the intracortical LFP beta band power (P2: F = 3.634, p = 0.027; P4: F = 0.982, p = 0.375), with BCI + 2-Back LFP beta band power (median: 26.02 dB, IQR: 25.33-26.59) being significantly lower than BCI Only (median: 26.22 dB, IQR: 25.48-26.72, p < 0.05). We examined raw beta band power during the reach phase (vs. normalized to rest) so an increase in power signifies less beta band desynchronization and therefore less movement-related modulation. Only P4 showed a significant difference in the intracortical firing rate (P2: F = 2.53, p = 0.081; P4: F = 4.542, p = 0.011), where firing rate was higher during BCI + 2-Back (median: 23.16 Hz, IQR: 21.79-24.83) compared to BCI Only (median: 23.10 Hz, IQR: 21.59-25.43, p < 0.05) or BCI + 1-Back (median: 23.00 Hz, IQR: 21.77-24.61, p < 0.05) signifying a stronger movement-related signal. While statistically significant, the absolute value of the change was quite small. Finally, neither participant exhibited differences in neural reaction time across conditions (P2: F = 2.48, p = 0.084; P4: F = 0.856, p = 0.425).

**Figure 5:**
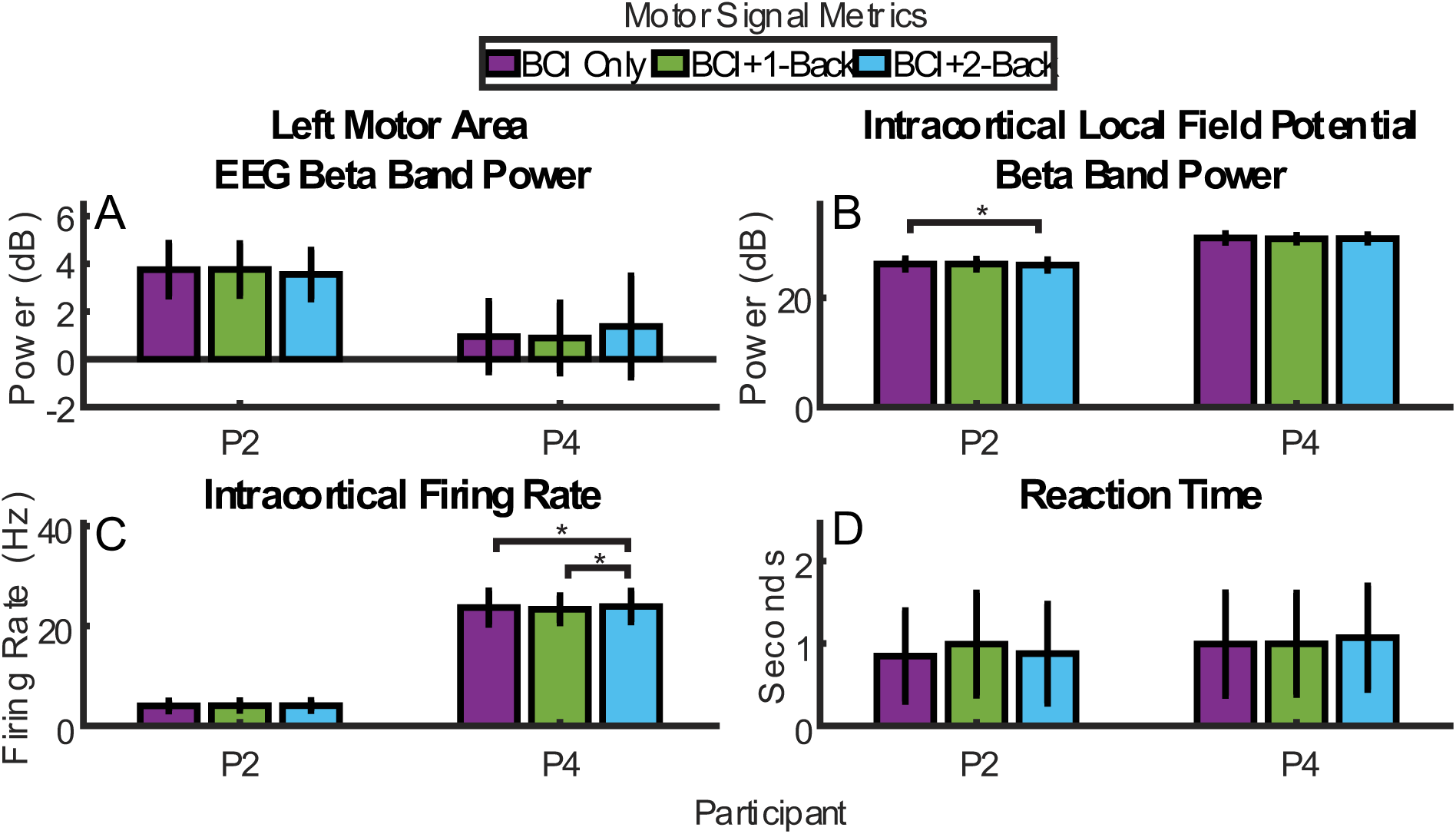
Measures of motor related activity from EEG and intracortical recordings, including A: EEG beta band power, B: Intracortical LFP beta band power, C Intracortical Firing Rate, D Neural Reaction Time. P2 exhibited a decrease in intracortical LFP beta band power in BCI + 2-Back compared to BCI only, indicating an increase in the motor signal. P4 exhibited an increase in intracortical firing rate in BCI+2-Back compared to BCI Only and BCI + 1-Back, signifying an increase in the motor signal.

## DISCUSSION

This study explored the effects of attentional load on iBCI performance and attention-and movement-related neural signals. The N-back task had little impact on iBCI performance even though it did cause changes in EEG correlates of attention. iBCI cursor control performance remained robust to attentional load even in a realistic and complex task involving click and drag actions. Movement-related neural activity measured intracortically and with EEG also remained stable during dual-tasking conditions when compared to BCI alone.

### Performance is Robust to Attentional Load

Participants exhibited few changes in performance across the different conditions, with only P2 displaying a slight increase in normalized path length in BCI+1-Back and a more moderate increase in total trial completion time as compared to BCI+2-Back. While such robustness is not unknown in the literature, the high level of attentional load here makes this nonetheless surprising. We built on previous iBCI work related to the impact of distractions (Guthrie et al., 2021) by utilizing a more complex task design (that included target-guided reaching and grasping to control a virtual computer mouse vs only simple left/right translational movement), expecting a greater impact of attentional load on performance. Our results support the idea that iBCI control is robust across multiple levels of attentional load.

The observed performance robustness is striking when comparing iBCI to EEG-BCI. Studies of the EEG BCI have found comparatively greater effects of attention. In one study, there was a large effect 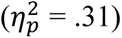 that led to a 20% drop in BCI success rate during a triple task of 1D BCI control, maintaining cruise altitude on a flight simulator, and performing attentional tasks (Vecchiato et al., 2016). In another study, EEG-BCI classification accuracy drops, in a BCI+speech task, ranged from 5% to 10% depending on the classifier used (İşcan & Nikulin, 2018). Emami & Chau, 2020, showed a range of performance of nearly 30% within low-performance BCI users as cognitive load increased. Foldes & Taylor, 2013, also show a decrease of up to 20% in BCI success rate during high cognitive load, along with slightly worse trial completion times and path efficiency. By contrast, our worst performing participant, P2, had a success rate drop of less than 10% on average. These results indicate that iBCI may be more robust against distractors than EEG-BCI. As such, the study of attentional load may be of greater importance to EEG-BCI researchers than those working with iBCI with the caveat that iBCI applications continue to become more complex and higher-dimensional (Collinger, Wodlinger, et al., 2013; Pandarinath et al., 2017) and may be more impacted by changes in attention as they do so.

One potential explanation for the lack of drop in performance is that participants ignored the N-Back in favor of the BCI task. However, we confirmed that the participants were engaged in the N-back task, both when performing the N-Back by itself and in conjunction with the BCI task, by measuring accuracy, which was above 80% in all N-Back conditions. This indicates a high level of engagement with the N-Back task, and that the consistent BCI performance is not due to ignoring the cognitive task during the dual task conditions. Participants subjectively reported that the dual tasking conditions required more mental effort (rated from 0-10) than the BCI only condition. For P2, both BCI+1-Back and BCI+2-Back required more mental effort compared to 1-Back only (Fig S2). For P4, BCI+2-Back required more effort than 1-Back only and BCI Only.

It is interesting to note that the participant whose performance did change slightly, P2, has been implanted for nearly ten years and has experienced an expected decline in signal quality (Sponheim et al., 2021) and overall BCI performance. It is possible that this more degraded performance and signal is more vulnerable to the dual-tasking compared to P4. This is supported by the findings of Emami & Chau, 2020, who found cognitive distractors to impact low-performing participants negatively but not high-performing participants. It is intuitive to assume lower-performing participants require more concentration to maintain performance.

### Correlates of Attention Change as Attentional Load Increases

In this study, we observed a complex relationship between EEG theta power and attentional load. In previous dual tasking studies, theta power has been found to increase compared to single tasking (Ozdemir et al., 2016). Although we also saw this effect, it was not consistent across conditions. P4 displayed an increase only in BCI+2-Back, the highest load condition. This could imply that BCI+1-Back was simply not complex enough to induce altered theta in P4 despite the subject reporting it required more mental effort than BCI Only. In the case of P2, the 1-back condition led to the highest theta band power with or without concurrent BCI control.

Alpha power exhibits a similar trend in P2 for both dual tasking and single tasking. However, this goes against our initial hypothesis of alpha decreasing during attentional load. Alpha and theta band EEG power both follow an inverted U-shaped trend in P2. Similar trends in neural metrics of cognitive load have been seen in single-tasking contexts. One study measuring prefrontal cortex activation in fMRI during an N-Back, ranging from N=1 to N=6, found that activation was highest at N=3 but then proceeded to decline (Lamichhane et al., 2020). One interpretation is that this reflects a shift in processing strategy where the brain regions that are active shift to handle the increasing load.

In the literature, alpha power often decreases during dual tasking and attentional tasks (Kahya et al., 2022). Our study observed an increase, but this is not unique. A previous study of mental workload during operation of a P300 EEG-BCI found increased alpha during dual tasking (Käthner et al., 2014). As in our study, their secondary cognitive tasks were presented through audio, and they noted that auditory stimulation has been found to increase alpha activity. They also discuss the alpha inhibition theory (Klimesch et al., 2007), which states alpha increases to inhibit non-essential brain regions to maintain task performance. İşcan & Nikulin also found that alpha power was negatively correlated with performance when BCI users had to listen to verbal counting (performance decreases with increasing alpha power), attributing the increase in power to attempts by participants to inhibit the distraction sounds. Overall, it is possible that the increased alpha in 1-Back is due to participants attempting to inhibit auditory-related stimuli so as to better handle increasing attentional load.

### Movement-Related Activity Remained Stable under Attentional Load

We initially hypothesized that the changes in performance and the motor signal would occur in response to increased attentional load, with the latter providing a potential mechanism for why attentional load impacts performance. Overall, the magnitude of changes observed in movement-related activity was very small. P2 had a slightly lower intracortical LFP Beta band power in the BCI + 2-Back condition. P4 had a slightly higher firing rate in the BCI + 2-Back condition. While the changes are very small, overall, this implies both participants exhibited a stronger motor signal in BCI + 2-Back, perhaps as a compensatory mechanism to maintain performance under higher load.

### Limitations

One limitation of this study is that we relied on EEG to measure attention. Although this is common in the literature, the potential for variability and noise may obscure some effects of attentional load. One of our participants did indeed have a good amount of noise that necessitated removing many trials. Another limitation is that participants were able to handle dual tasking with great performance. We used tasks that were representative of typical BCI use and are generally more difficult than simple center out cursor tasks, but it seems they still may not have been enough to push attentional load to the fullest. Future studies should ensure the tasks are sufficiently challenging to fully sample the range of possible attentional load conditions. Finally, this study is limited to 2 participants due to the nature of intracortical BCI trials, which inherently have small sample sizes. The data provided here suggest that factors such as signal quality and baseline BCI performance ability may affect the impact of attentional load, but additional testing is warranted to clearly delineate those effects.

## CONCLUSION

This work adds to the small literature investigating the impact of attentional load on BCI performance. It is one of very few studies that have looked at the impact of attention or distraction during iBCI control (Guthrie et al., 2021; Stavisky et al., 2020). Unlike these previous studies, neural correlates of attention were quantified with whole brain EEG measures. The N-back was sufficient to drive changes in attentional load, but dual tasking with BCI resulted in very minimal changes in performance. iBCI cursor control was robust to changes in attentional load, but more complex applications may be more susceptible to attention-related neural changes.

## Data Availability

Data are available upon request at http://doi.org/10.18120/ar71-yw82.

## Supplemental Material

**Figure S1:**
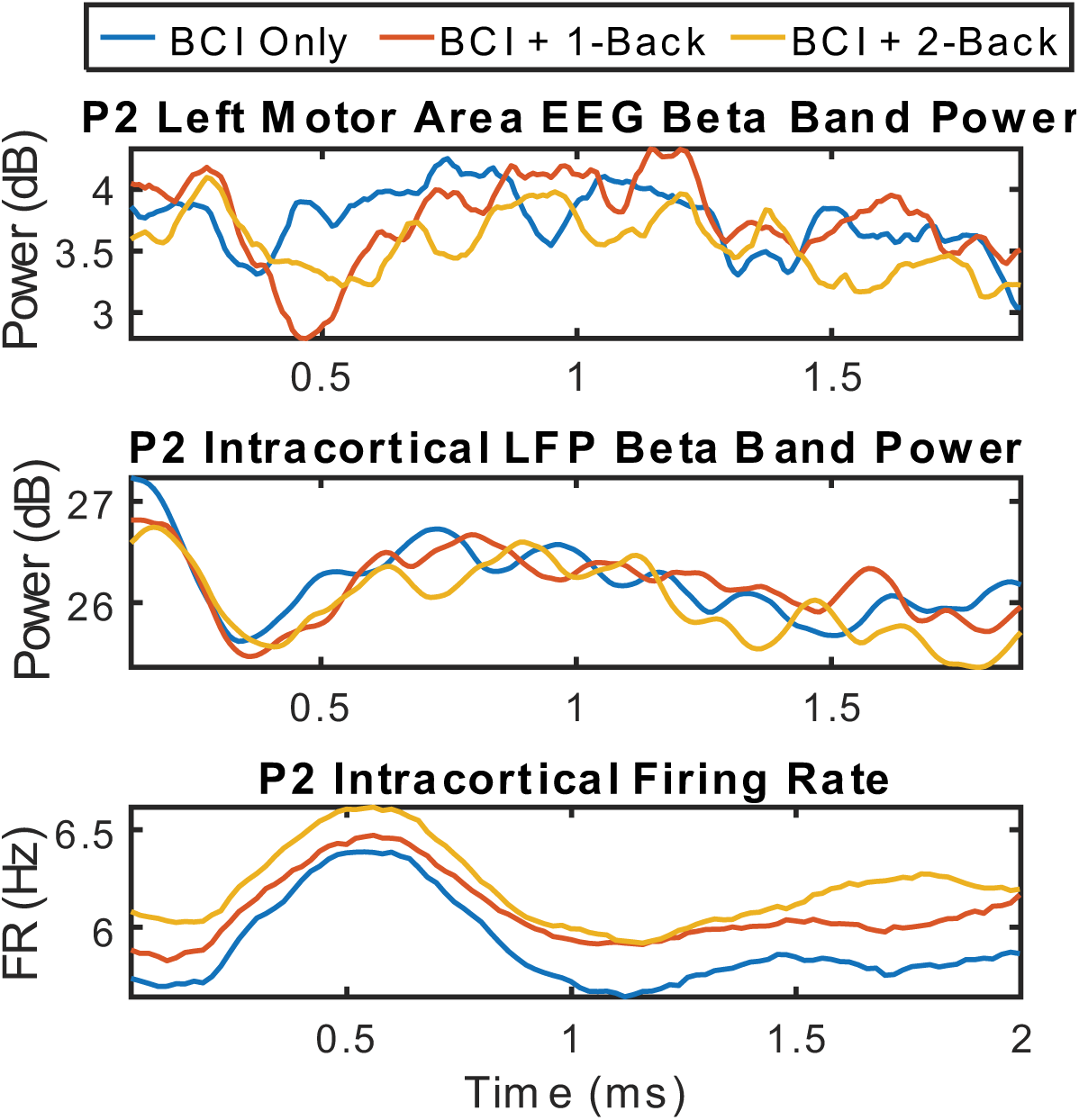
Average motor signal for P2 as measured using EEG beta band power (top), intracortical local field potential beta band power (middle), and intracortical firing rate (bottom). Data is averaged across channels within a region and then across all trials.

**Figure S2:**
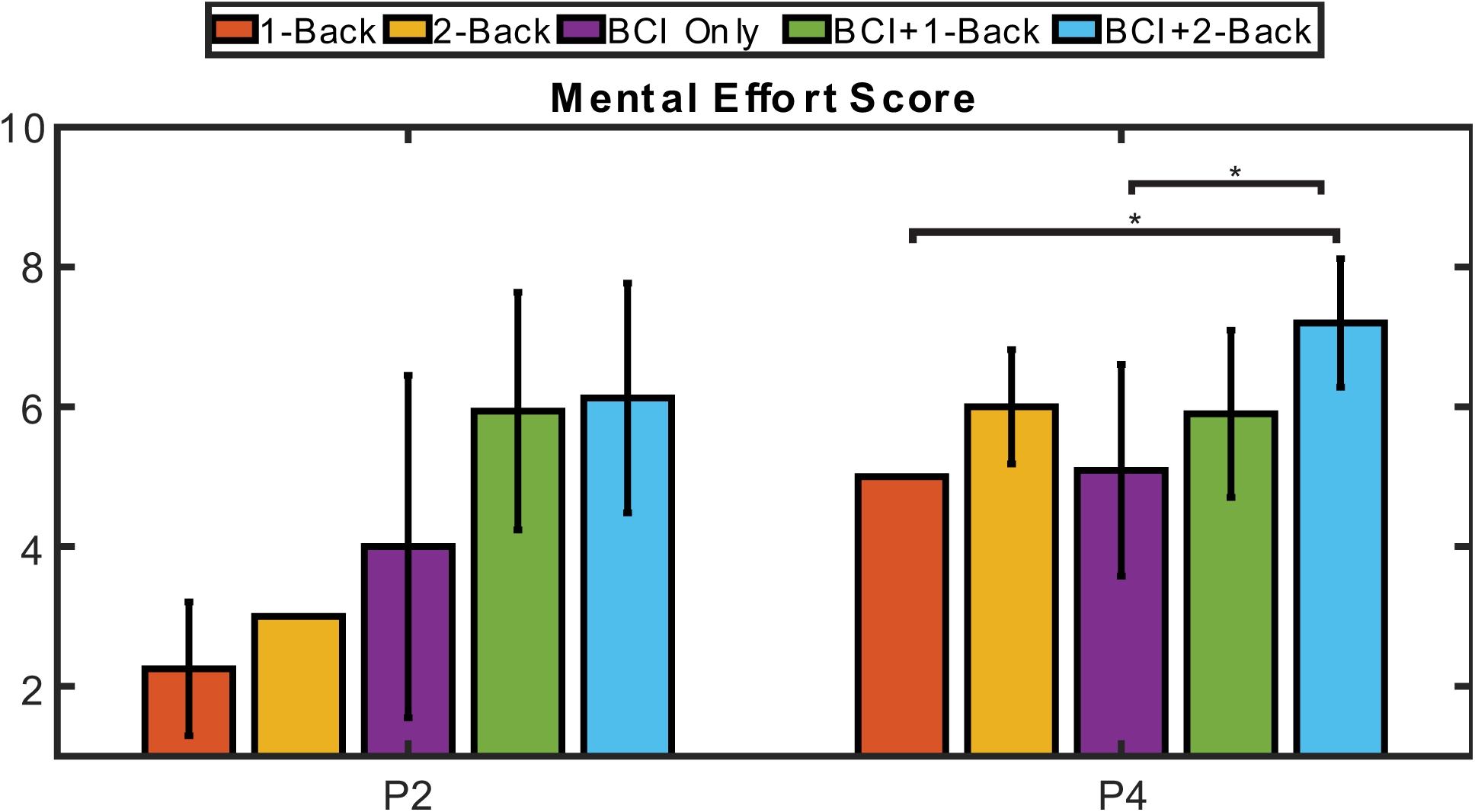
Mental effort scores for each condition and participant. Scores increase during higher load levels or during dual tasking. Using a regression model as described in methods, we observed a significant difference in mental effort across conditions. P2’s model is significant but individual comparisons do not survive post-hoc comparisons (F = 3.85, p = 0.02). P4 does display significant differences (F = 6.98, p = 0.0005), with BCI + 2-Back having a higher average mental effort score compared to 1-Back and BCI Only.

## Acknowledgements

We would like to acknowledge Dr. Matt Smith for his helpful input and discussion, as well as Debbie Harrington for her support in regulatory management, Carleigh May and Gracie Hilber for their aid in data collection, and Dr. Michael Boninger for serving as sponsor-investigator of the clinical trial in this study. We would also like to thank the study participants for their invaluable contributions to this study.

